# Impact of membrane lipid polyunsaturation on dopamine D2 receptor ligand binding and signaling

**DOI:** 10.1101/2022.01.04.474945

**Authors:** Marie-Lise Jobin, Véronique De Smedt-Peyrusse, Fabien Ducrocq, Asma Oummadi, Rim Baccouch, Maria Hauge Pedersen, Brian Medel-Lacruz, Pierre Van Delft, Laetitia Fouillen, Sébastien Mongrand, Jana Selent, Tarson Tolentino-Cortez, Gabriel Barreda-Gómez, Stéphane Grégoire, Elodie Masson, Thierry Durroux, Jonathan A. Javitch, Ramon Guixà-González, Isabel D. Alves, Pierre Trifilieff

## Abstract

The heterogenous and dynamic constitution of the membrane fine-tunes signal transduction. In particular, the polyunsaturated fatty acid (PUFA) tails of phospholipids influence the biophysical properties of the membrane, production of second messengers, or membrane partitioning. Few evidence mostly originating from studies of rhodopsin suggest that PUFAs directly modulate the conformational dynamic of transmembrane proteins. However, whether such properties translate to other G protein-coupled receptors remains unclear. We focused on the dopamine D2 receptor (D2R), a main target of antipsychotics. Membrane enrichment in n-3, but not n-6, PUFAs potentiates ligand binding. Molecular dynamics simulations show that the D2R preferentially interacts with n-3 over n-6 PUFAs. Furthermore, even though this mildly affects signalling in heterologous systems, *in vivo* n-3 PUFA deficiency blunts the effects of D2R ligands. These results suggest that n-3 PUFAs act as allosteric modulators of the D2R and provide a putative mechanism for their potentiating effect on antipsychotic efficacy.

## Introduction

Biological membranes are not homogeneous bilayers but rather composed of different lipid species with various chemical and biophysical properties that actively modulate protein localization and function, signaling or vesicular trafficking ^1^. However, several aspects of membrane complexity including the impact of lipid heterogeneity across tissues, cells, and subcellular compartments on cell signaling and physiology are only starting to emerge.

This is particularly true for neuronal function for which the impact of membrane lipid composition has been largely overlooked, despite that the brain has the second highest lipid content after adipose tissue ^2^. Yet, convergent findings support some links between lipid biostatus and mental health. For instance, a decrease in the levels of n-3 polyunsaturated fatty acids (PUFAs)-containing phospholipids has been consistently described in a subset of patients suffering from neurodevelopmental psychiatric diseases ^3^. Likewise, recent transcriptomic profiling studies revealed a common pattern in fatty acid metabolic pathways in several psychiatric disorders ^4,5^. Moreover, n-3 PUFA deficiency in rodent models has been associated with putative pathophysiological mechanisms involving alteration in neurogenesis, neuronal migration, neuromodulation, and neuroinflammatory processes in various brain regions ^2,6,7^. These findings remain largely correlative and the precise mechanisms by which PUFA biostatus directly or indirectly accounts for changes in neuronal function remain largely unknown. Nevertheless, several studies have demonstrated that membrane lipids regulate the function of key transmembrane proteins ^8^ including ion channels and GPCRs ^9,10^.

The activity of the dopamine D2 receptor (D2R) is particularly relevant in this context. Various psychiatric disorders, including those where levels of brain n-3 PUFAs are low ^11^, display altered patterns of D2R-dependent signaling, and this GPCR is, therefore, a key target in several pharmacological treatments. Similarly, different studies in rodent models show that n-3 PUFA deficiency affects dopamine transmission and related behaviors ^12,13^. In line with this evidence, we recently reported a unique vulnerability of D2R-expressing neurons to PUFA biostatus that directly accounts for the motivational deficits induced by n-3 PUFA deficiency ^14^. Early experimental studies have reported that n-3 PUFAs, namely docosahexaenoic acid (DHA, 22:6), enhance the function of the prototypical A GPCR rhodopsin ^15–17^. Subsequent studies showed that DHA-containing phospholipids preferentially solvate rhodopsin ^18,19^, which could impact receptor function. In addition, recent *in silico* studies ^20,21^ demonstrate that the D2R also displays a preference for DHA interaction that could modulate receptor partitioning into specific membrane signaling platforms, as shown by early experiments ^22^. While these findings suggest that membrane PUFA composition could influence the activity of the D2R, there is still no direct evidence of such an effect.

In this work, using both cell membrane extracts and model membranes, we found that DHA, but not n-6 PUFA docosapentaenoic acid (DPA, 22:5), enhances D2R ligand binding affinity. Strikingly, while DPA and DHA only differ in one double bond, molecular dynamics (MD) simulations show that they have a completely different propensity for interaction with the D2R. Interestingly, membrane enrichment in either DHA or DPA have no effect on agonist-induced, G_i/o_ protein-dependent inhibition of cAMP production. However, both DHA and DPA similarly enhance the maximal efficacy of the D2R to recruit β-arrestin2 upon agonist stimulation. Finally, we provide *in vivo* evidence that DHA deficiency blunts the behavioral effects of D2R ligands. Altogether, these data highlight the impact of membrane PUFA composition on D2R activity and suggest that DHA acts as an allosteric modulator of the receptor.

## Results

### 1. Membrane n-3 PUFAs, but not n-6 PUFAs, modulate the binding affinity of D2R ligands

To determine the impact of membrane lipid unsaturation on D2R ligand binding, HEK cells were incubated with PUFAs, which is known to result in their esterification in *sn*-2 position of phospholipids ^23^. We used two different PUFAs, the n-3 PUFA DHA (C22:6) and the n-6 PUFA DPA (C22:5), which display the same carbon chain length, but differ in one double bond. A lipid analysis confirmed the enrichment of phospholipids in DHA or DPA and monitored its efficient incorporation in cell membranes, leading to comparable proportion of either PUFA after treatment (increase of 19% for DHA and 20% for DPA) (Fig. S1). We measured the binding affinity of D2R ligands on supported cell membranes expressing the D2R using plasmon waveguide resonance (PWR) spectroscopy ^24^. The resonance position shifts were followed in *p*- (perpendicular to the sensor surface) and *s*-polarisation (parallel to the sensor surface) upon poly-L-Lysine (PLL) coating of the PWR sensor and subsequent cell membrane adhesion and detachment (Fig. S2A and B). The cell membrane fragment deposition led to positive shifts similar to those observed for synthetic lipid bilayer formation ^25^. We measured the resonance shifts upon incremental addition of ligand when the time curve reached a plateau (Fig. S2C and D) and led to shifts in the minimum resonance. These cumulative shifts were plotted as a binding curve which allowed calculating a dissociation constant (K_D_) and measuring a plateau of the resonance shifts (Fig. S3). The K_D_ calculated for the agonist quinpirole and the antagonist spiperone were comparable to the ones reported in the literature, validating our system and confirming that the receptor maintains functionality in these conditions (Fig. S3C) ^26,27^. The results from these experiments show that membrane DHA enrichment enhances ligand binding affinities for agonists (quinpirole and dopamine), antagonist (spiperone), and partial agonist (aripiprazole) (Fig. 1). However, DPA slightly decreased D2R binding affinity for quinpirole and spiperone (Fig. 1A and E) and had no effect for dopamine and aripiprazole (Fig. 1C and G). We further confirmed the potentiating effect of membrane DHA enrichment on D2R ligand binding affinity for the spiperone-derived fluorescent ligand NAPS-d2 through fluorescence anisotropy ^28,29^ (Fig. S4) and in partially purified D2R reconstituted in model membranes (Fig. S5).

**Figure 1.**
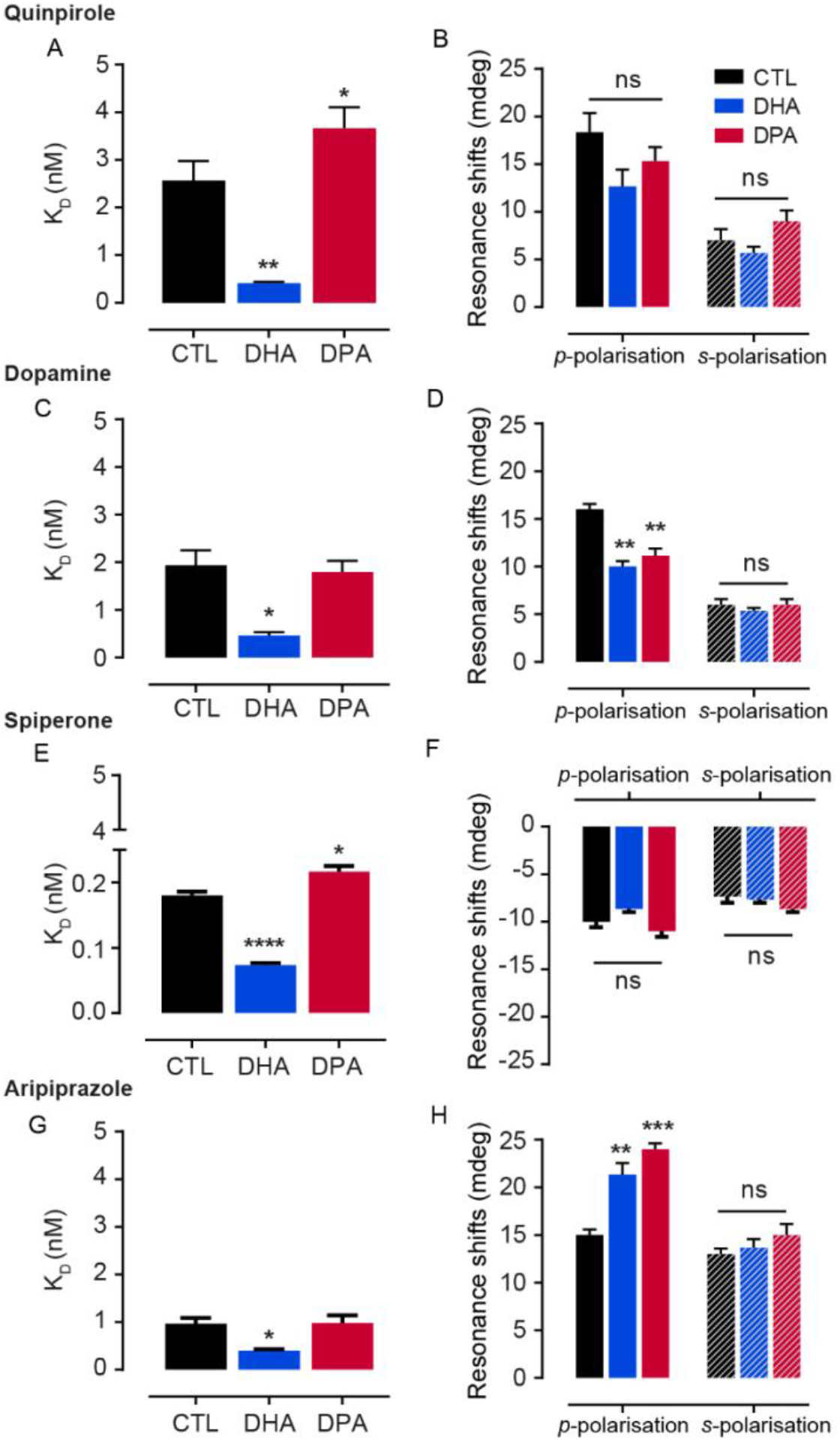
D2R binding affinity in PUFA-enriched cell membranes shown by PWR. PUFA effect on binding affinity and receptor conformational changes, respectively, for quinpirole (A and B), dopamine (C and D), spiperone (E and F) and aripiprazole (G and H). Data are mean ± SEM values from 3 independent experiments with **** p<0.0001, *** p <0.001, ** p<0.01, * p<0.05 by two-tailed unpaired t-test.

To study ligand-induced receptor conformational changes, we quantified the direction and magnitude of minimum resonance in *p*- and *s*-polarizations, which account for changes along the receptor long and short axis, respectively. Resonance shifts obtained upon ligand addition were positive in both polarisations for agonists (quinpirole, dopamine) and partial agonist (aripiprazole), and negative for the antagonist spiperone (Fig. 1F). These results show that different classes of ligands induce distinct conformational changes upon binding to the D2R, as previously reported for other GPCRs ^30–34^. While the presence of membrane PUFAs did not significantly influence the changes in conformation induced by quinpirole and spiperone (Fig. 1B and F), they impacted the conformation of dopamine- and aripiprazole-bound D2Rs regarding the anisotropy of the conformational state that became less anisotropic for dopamine and more anisotropic for aripiprazole (as shown by the respective decrease and increase in *p*-polarization shifts; Figs. 1D and 1H).

### 2. MD simulations reveal that n-3 and n-6 PUFAs differentially interact with the D2R

With the aim of gaining structural insights into the differential modulation exerted by DPA and DHA in PUFA-enriched cell membranes, we used MD simulations to study the interaction between the D2R and membranes enriched in either DHA or DPA. We first simulated four replicas of the D2R embedded in a multicomponent membrane (see Methods) enriched in either DHA-containing phospholipids (1-stearoyl-2-docosahexaenoyl-sn-glycero-3-phosphocholine, SD(h)PC) or DPA-containing phospholipids (1-stearoyl-2-docosa(p)entaenoyl-sn-glycero-3-phosphocholine, SD(p)PC). In line with previous reports, ^20,21^, SD(h)PC tends to displace saturated phospholipids from the membrane “solvation” shell that surrounds the D2R, due to the much higher propensity of DHA tails for the interaction with the receptor (Fig. 2A). Strikingly, the loss of just one double bond (i.e. replacing DHA by DPA) is enough to abolish this propensity, as shown by the much lower interaction ratio of SD(p)PC molecules with the receptor during the simulation (Fig. 2B). Specifically, the propensity for the interaction with the D2R is three-fold higher for DHA tails (Table 1).

**Figure 2.**
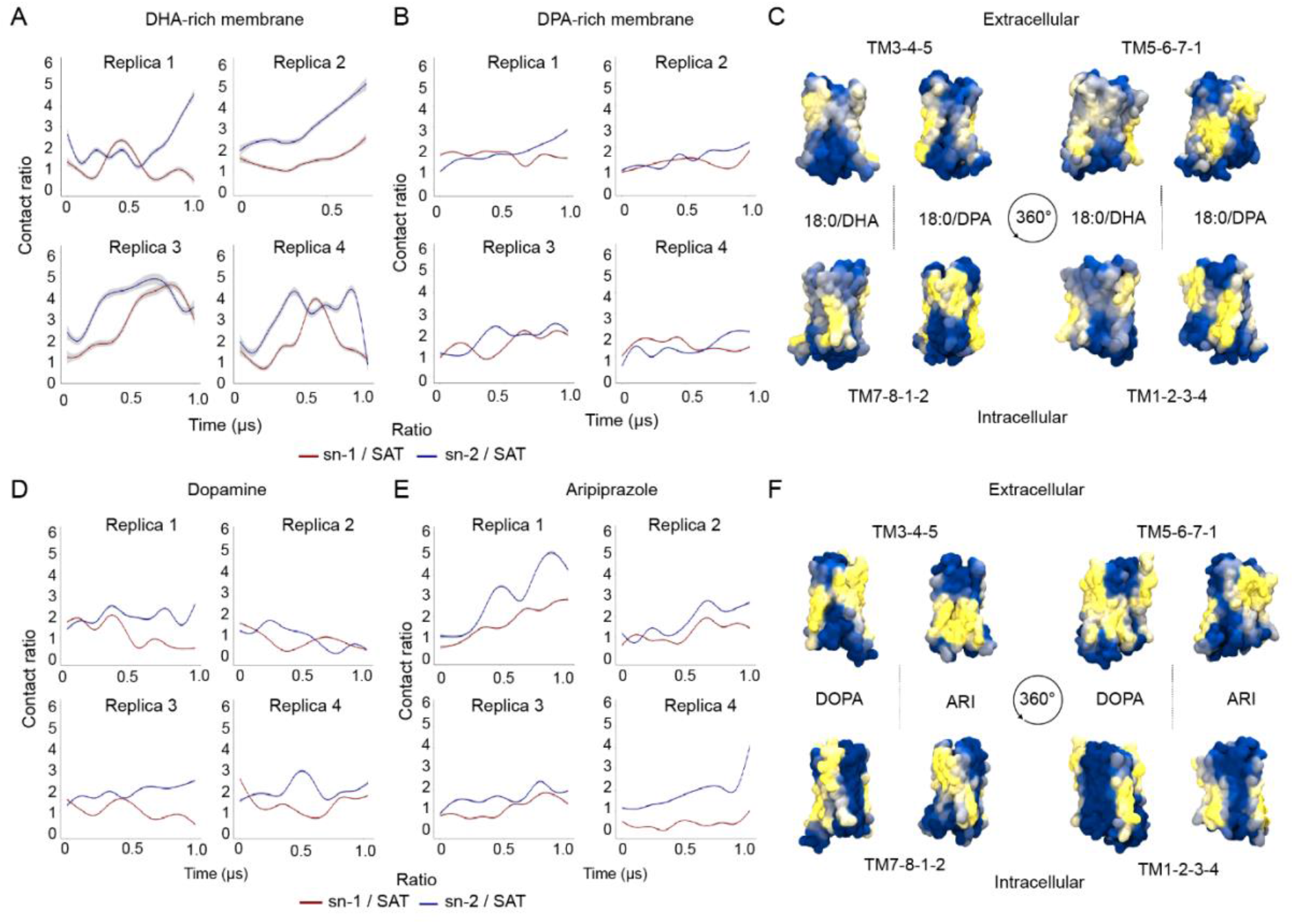
Lipid-protein contacts during MD simulations. **(A, B, D, E)** Relative proportion of atomic lipid-protein contacts (y-axis) over time (x-axis) for the simulations of the apo state of the D2R embedded in DHA- (A) versus DPA-rich (B) membranes, and the simulations of dopamine- (D) versus aripiprazole-bound D2R (E) embedded in DHA-rich membranes. Specifically, figures depict the contact ratio of each SD(h)PC or SD(p)PC chain versus all saturated lipids (i.e. DPPC, DSPC, and PSM) in the system (i.e. sn-1 / SAT and sn-2 / SAT) is depicted (see Methods for a detail description of these ratios). **(C, F)** Average lipid occupancy map for the simulations of the apo state of the D2R embedded in DPA-versus DHA-containing phospholipids (C), and the simulations of dopamine- (DOPA) versus aripiprazole (ARI)- bound D2R embedded in DHA-rich membranes (F). The percentage of frames where SD(h)PC or SD(p)PC lipid molecules contacted the D2R is depicted in a blue (low) to yellow (high) color gradient mapped to the surface of the receptor.

**Table 1.** Evolution of lipid-protein contacts during MD simulations. The table shows the average relative proportion of lipid-protein contacts at four stages of the simulation. Contact ratios represent the number of atomic contacts between each SD(h)PC or SD(p)PC chain (i.e. sn-1 or sn-2) and the D2R, with respect to the number of contacts between saturated phospholipids (i.e. DPPC, DSPC, and PSM) and the protein (see Methods for more details).

| Ratio | 0 - 250 ns | 250 - 500 ns | 500 – 750 ns | 750 ns - 1 μs |
| --- | --- | --- | --- | --- |
| SD(h)PC<sub>sn-1</sub> | 0.97 | 1.69 | 2.95 | 2.42 |
| **SD(h)PC<sub>sn-2</sub> (DHA)** | **2.01** | **3.05** | **4.61** | **4.08** |
| SD(p)PC<sub>sn-1</sub> | 1.49 | 1.48 | 1.48 | 1.63 |
| **SD(p)PC<sub>sn-2</sub> (DPA)** | **1.47** | **1.95** | **2.28** | **2.33** |

From these results, it is reasonable to speculate that the more flexible lipid corona built around the D2R in DHA-rich membranes could allow for greater dynamics within the receptor, which could potentially enhance protein conformational selection during the ligand binding process. This scenario would accord with our ligand binding experiments, where DHA-rich membranes consistently induced lower K_Ds_, i.e. higher binding affinities, than DPA-rich membranes (Fig. 1). Moreover, our simulations show that despite DHA and DPA only differing in one double bond, they display a different pattern of interaction with the receptor (Fig. 2C). Some of these differences could also impact ligand binding affinity at the D2R. For example, SD(p)PC lipid molecules display a clear preference over SD(h)PC for interaction with the extracellular segments of transmembrane helix (TM) 1, TM2, and TM7, which are known to define entry crevices for water and phospholipid headgroups into the binding pocket ^35^ (Fig. 2C, bottom left).

We also simulated the D2R bound to dopamine or aripiprazole molecules, and embedded into a multicomponent membrane enriched in SD(h)PC (see Methods). Interestingly, while the aripiprazole-bound D2R simulations also showed an increased level of DHA around the receptor (Fig. 2E), the DHA solvation effect was absent or even diminished in dopamine-bound D2R simulations (Fig. 2D). Specifically, the propensity for the interaction between DHA tails and the protein was approximately two-fold higher in the aripiprazole-bound D2R simulations (Table 2).

**Table 2.** Evolution of lipid-protein contacts during ligand-bound D2R simulations. The table shows the average relative proportion of lipid-protein contacts at four stages of the simulation. Contact ratios represent the number of atomic contacts between each SD(h)PC chain (i.e. sn-1 or sn-2) and the dopamine (DOPA)- or aripiprazole (ARI)-bound D2R, with respect to all saturated lipids (i.e. DPPC, DSPC and PSM) (see Methods for exact membrane composition and details on the calculation of these ratios).

| | Ratio | 0 - 250 ns | 250 - 500 ns | 500 - 750 ns | 750 ns - 1 μs |
| --- | --- | --- | --- | --- | --- |
| DOPA | SD(h)PC<sub>sn-1</sub> | 1.16 | 0.98 | 1.00 | 0.81 |
|  | **SD(h)PC<sub>sn-2</sub> (DHA)** | **1.78** | **1.63** | **1.62** | **1.73** |
| ARI | SD(p)PC<sub>sn-1</sub> | 0.97 | 1.46 | 1.65 | 1.67 |
|  | **SD(p)PC<sub>sn-2</sub> (DHA)** | **1.75** | **2.87** | **3.12** | **3.07** |

Furthermore, as shown in Fig. 2F, the presence of the ligand in the binding pocket induced a different pattern of interactions between SD(h)PC and the D2R, including a different interaction signature at the extracellular segment of the receptor where ligand binds (Fig. 2F, upper left and right panels)

### 3. n-3 and n-6 PUFAs do not influence the D_2_R G_αi/o_-mediated signaling pathway but enhance maximal recruitment of β-arrestin2

In order to evaluate the downstream effect of PUFAs, we performed dose-response experiments where we probed the signaling efficacy of the D2R in the presence and absence of PUFAs. First, we treated cells with 10 μM PUFAs (Fig. S6) and measured the inhibition of forskolin-induced cAMP production upon quinpirole, dopamine and aripiprazole binding as a proxy for G_i/o_ protein signaling. As shown in Fig. S7A and Table S1, both control and PUFA-rich cells display a similar agonist-dependent decrease in cAMP production, which suggests that PUFAs do not modulate the G_i/o_ protein signaling pathway. Next, we used the same experimental conditions to study the G_i/o_ protein-independent signaling pathway by measuring β-arrestin2 recruitment. In these experiments, while PUFAs did not affect the potency of β-arrestin2 recruitment (i.e. EC_50_), they either substantially (aripiprazole) or slightly (quinpirole and dopamine) affected its efficacy (i.e. E_max_) (Fig. S7C, S7D, S7E). To confirm this trend, we treated cells with a higher amount of PUFAs (i.e. 30 μM) (Fig. S6) and we did not observe any impact of PUFAs enrichment on basal cAMP production (Fig. S7B). We then performed dose-response experiments upon ligand stimulation. Higher PUFA manipulations did not influence cAMP production or the EC_50_ for β-arrestin2 recruitment (Figs. 3A, 3C, and 3E), but significantly increased the E_max_ of β-arrestin2 recruitment for all ligands in both DHA- and DPA-rich cells (Figs. 3B, 3D, and 3F).

**Figure 3.**
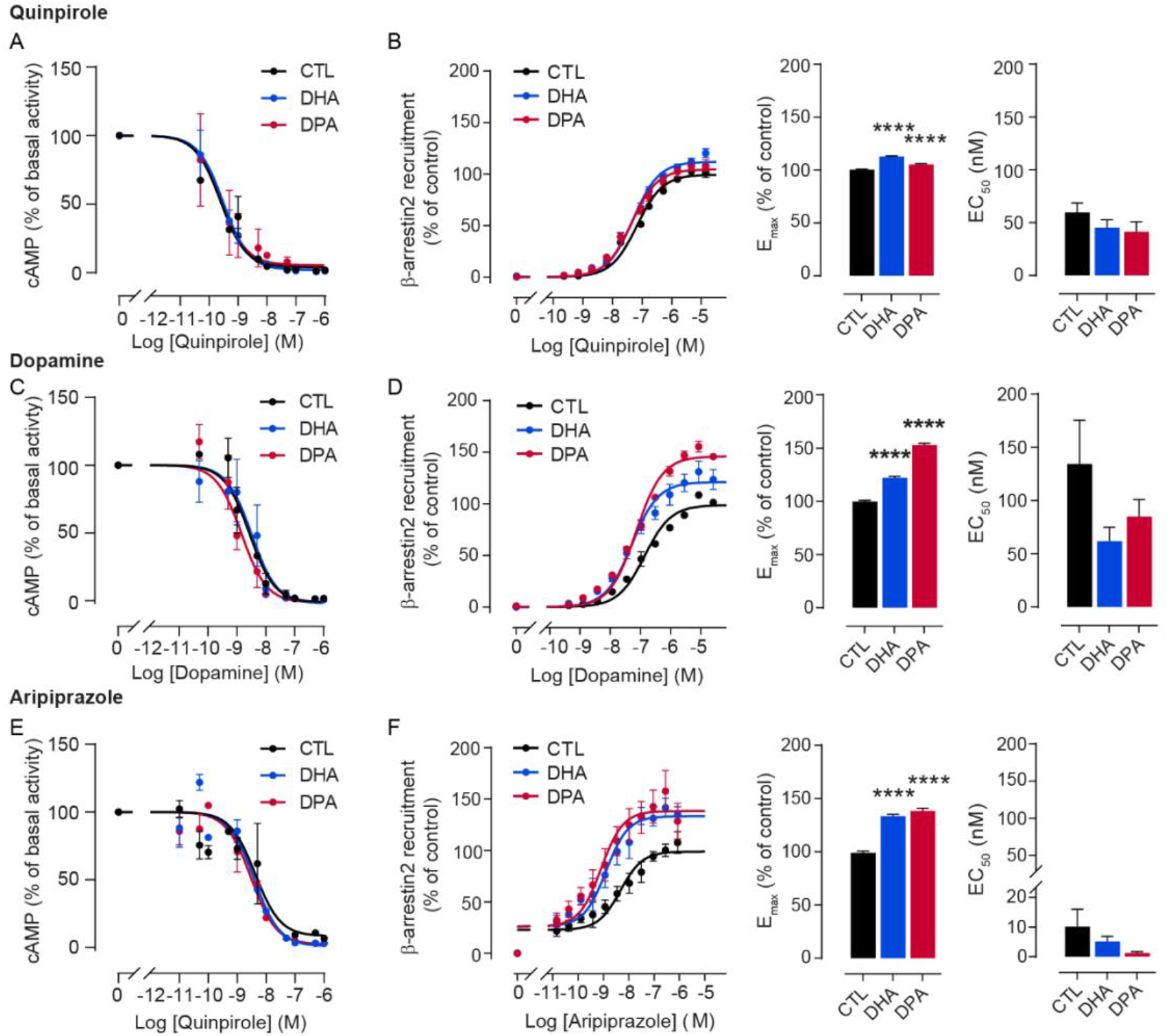
Forskolin-stimulated cAMP production and β-arrestin2 recruitment upon ligand stimulation and with different membrane PUFAs enrichment. **(A, C, E)** Dose-response experiments of cAMP production with the D2R ligands quinpirole (A), dopamine (C) and aripiprazole (E), on forskolin-stimulated cells incubated in the presence of 0.03% ethanol as control, 30 μM DHA and 30 μM DPA n-6. Values are expressed as the percentage of cAMP in the absence of agonist in n=3 independent experiments. **(B, D, F)** Quinpirole (B), Dopamine (D) and aripiprazole (F) activity on D2R mediated β-arrestin2 assay in CHO-K1 cells expressing the DRD2L (left panel), and associated Emax (middle) and EC50 (right panel) under 30 μM PUFAs enrichment. Data are mean ± SEM values from three independent experiments with **** p<0.0001, *** p <0.001, * p<0.05 by two-tailed unpaired t-test.

Altogether, our data reveal that both DPA and DHA exert a similar effect on downstream signaling, namely they do not affect G_i/o_ protein activation but they are able to enhance β-arrestin2 recruitment efficacy.

### 4. n-3 PUFA deficiency alters D2R ligand-induced alterations in locomotion and motivation

To translate our *in vitro* findings to *in vivo* settings, we used a validated model of n-3 PUFA deficiency in mice. Using such an approach, we previously demonstrated that i) n-3 PUFA deficiency leads to decreased and increased levels of DHA and DPA in membrane phospholipids, respectively and ii) n-3 PUFA deficient animals display motivational deficits that are due to impaired functionality of D2R-expressing neurons ^14^. Herein, using striatal extracts – in which D2R is highly expressed – we found that i) Gi protein recruitment at the receptor under quinpirole application is unaffected (Fig. 4A) and ii) phosphorylation of the glycogen synthase kinase β (GSK3β), a key signaling enzyme downstream of β-arrestin2 ^36^, is decreased at basal states in n-3 PUFA deficient animals (Fig. 4B and S8). These latter findings are consistent with our in vitro data showing that membrane PUFA enrichment potentiates the recruitment of β-arrestin2 at the D2R. Next, we used n-3 PUFA deficient animals to study behavioral responses to D2R ligands. Our results show that n-3 PUFA deficient animals display decreased locomotor response under quinpirole administration (Fig. 4C). Finally, we found that peripheral administration of aripiprazole blunted performance in an operant conditioning-based motivational task in control animals, while n-3 PUFA deficient mice were partially insensitive to aripiprazole (Fig. 4D).

**Figure 4.**
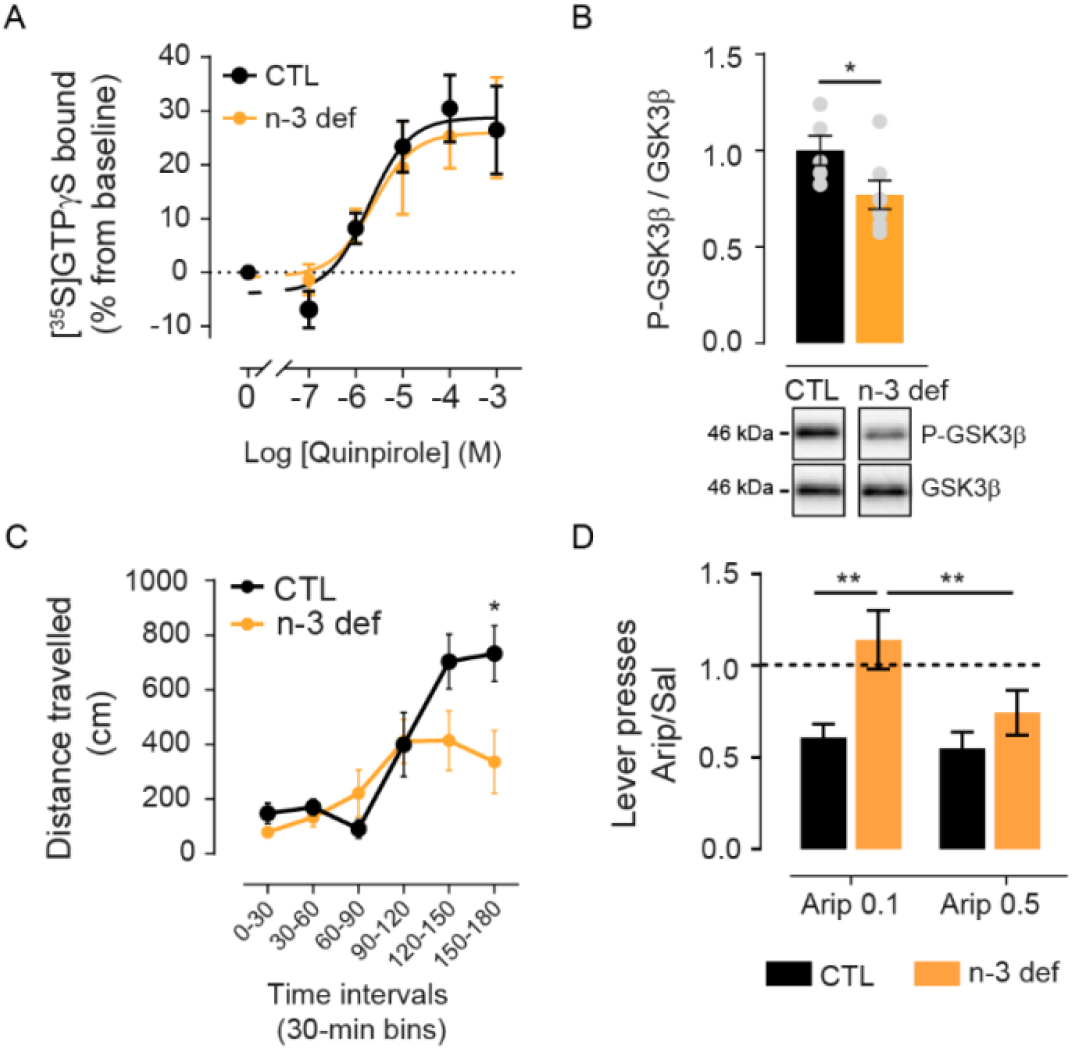
Effect of n-3 PUFA deficiency on D2R signalling and associated behaviors. **(A)** [^35^S]GTPγS assay on nucleus *accumbens* extracts upon increasing concentrations of quinpirole from control (CTL) or n-3 PUFA deficient animals (n-3 def). CTL : logEC50 = −5.22 ± 0.28 %, Emax = 30.06 ± 3.47 %; n-3 def : logEC50 = −5.06 ± 0.29; Emax = 28.15 ± 3.51 % **(B)** Western blots measuring the phosphorylation of GSK3β (P-GSK3β) relative to GSK3-β. Bars represent the mean of 5 and 7 subjects for CTL and n-3 def respectively and error bars the SEM. *p<0.05 by Mann Whitney test. **(C)** Locomotor response to 1 mg/kg quinpirole administration is represented as the distance travelled in cm divided in 30 min intervals in control (CTL, n=10) or n-3 def (n=10) animals. **(D)** Ratio of lever presses performed by animals under aripiprazole (0.1 or 0.5 mg/kg) over saline administration in a progressive ratio task in CTL (n=11) or n-3 def (n=10) animals. Data are mean ± SEM values. ** p< 0.01, * p<0.05 by two-way (RM) ANOVA test with a post-hoc Bonferroni test.

## Discussion

While most early reports on the preferential interaction between PUFAs and GPCRs involved n-3 PUFAs (i.e. DHA) and rhodopsin ^15–19^, recent studies have confirmed that DHA can also modulate the activity and function of other GPCRs including the cannabinoid CB1 receptor ^37^, the D2R ^20,21^ and the adenosine A2A receptor ^20,21,38^. Here, we focus on D2R interplay with PUFAs and show that (a) DHA-but not DPA-rich phospholipids potentiate the binding affinity of both agonists and antagonists for the D2R; (b) both DPA and DHA enhance β-arrestin2 recruitment efficacy without affecting G_i/o_ protein activity; and (c) n-3 deficient animal models display altered behavioral responses upon treatment with D2R ligands.

PUFA-containing phospholipids can have a strong influence on the physical and mechanical properties of biological membranes including thickness, bending or rigidity ^39^ and, hence, subtle changes in these properties can indirectly modulate the function of transmembrane proteins ^39–41^. Therefore, PUFA-rich membranes could potentially favor specific conformations of the D2R to modulate its ligand binding affinity. In particular, n-3 PUFAs provide transmembrane proteins with a uniquely flexible environment that enables such modulation ^41–43^. Interestingly, despite differing in just one double bond, n-6 PUFAs do not seem to increase membrane elasticity to the same extent ^44^. Furthermore, by increasing membrane fluidity and/or packing defects ^8^, PUFAs could change D2R accessibility for ligands thus modulating their affinity. In fact, most antipsychotics (i.e. D2R ligands) display high membrane partitioning properties ^45,46^. Of note, Lolicato et al. have recently shown that the entry pathway of dopamine into the D2R likely requires this neurotransmitter to first partition into the membrane ^47^.

On the other hand, different studies demonstrate that PUFAs tend to preferentially interact with GPCRs ^18,19,48,49^; thus, changes in ligand binding affinity could also be the result of direct interactions between PUFAs and the D2R. In the current study, MD simulations confirm that DHA tails preferentially interact with the D2R when compared to saturated ones, as previously reported ^20,21^. Intriguingly, our simulations show that this preferential interaction is lower or completely absent in the presence of aripiprazole or dopamine in the D2R binding pocket, respectively. This result suggests a more complex scenario where the conformational state induced by the ligand could also modulate the interaction propensity between the membrane and the receptor, as recently suggested for cholesterol and the oxytocin receptor ^50^. Interestingly, our simulations also show that n-6 PUFAs (i.e. DPA) have a much lower tendency to interact with the receptor, despite DHA and DPA only differing in one double bond. While further work is needed to unravel the precise molecular mechanisms behind the effect of DHA on ligand binding affinity, our work suggests that DHA can act as an allosteric modulator of the D2R, both by influencing the bulk membrane properties and by establishing direct interactions with the receptor.

However, our dose-response experiments in cells show that the enhancing effect of PUFAs on D2R ligand binding affinity does not correlate with an increased signal of relevant downstream protein effectors. On the one hand, enriching membranes with PUFAs did not alter cAMP production (i.e. G_i/o_ protein signaling). This result is in line with different early ^51,52^ and more recent reports ^53^. On the other hand, PUFA-rich membranes do enhance the maximal efficacy of β-arrestin2 recruitment, which seems to support a mechanism in which these lipids modulate specific protein effectors. This is compatible with the fact that PUFAs can alter the lateral organization of membranes 2^3,54–56^ or change the domain partitioning properties of GPCRs including the D2R ^20,21^. Changes in the properties of membrane microdomains could also affect the affinity of palmitoyl moieties for these domains, which could in turn affect receptor partitioning. As previously suggested ^57–62^, altering receptor partitioning could indirectly modulate receptor phosphorylation and, hence, influence β-arrestin coupling. It is also worth speculating that PUFA-induced membrane packing defects ^8^ could alter β-arrestin recruitment by modulating the membrane anchoring of C-edge loops ^63^. Therefore, one plausible albeit speculative hypothesis is that PUFAs could selectively modulate D2R signaling properties by altering membrane lateral organization and/or influencing β-arrestin anchoring to the membrane. It is worth noting that (a) the potency (EC_50_) of β-arrestin recruitment does not change, and (b) both DPA and DHA exert comparable effects on protein signaling whereas only DHA is capable of enhancing ligand binding. Altogether, these data suggest that the effects of PUFAs on D2R signaling are at least partially uncoupled from the changes in receptor binding properties. However, it should be noted that the relatively small changes in ligand binding affinity induced by PUFAs on ligand binding affinity might be difficult to detect using signalling assays in heterologous system. Nonetheless, such subtle modifications might have important physiological impact in the context of chronic alterations in PUFA levels, in particular when occurring during brain development.

This hypothesis is supported by our *in vivo* findings in a model of chronic deficiency in n-3 PUFAs ^14^. In fact, in accordance with our *in vitro* results, while n-3 PUFA deficiency does not seem to affect D2R binding of the G_i/o_ protein in the striatum, the decrease in phospho-GSK3 expression is consistent with an impairment of β-arrestin2-dependent signaling, since GSK3 phosphorylation depends on β-arrestin2 recruitment at the D2R ^36^. Notably, D2R-mediated locomotor response has recently been shown to also depend on β-arrestin2-, but not G_i/o_ protein-, mediated signalling ^64^. In line with this, we find that D2R agonist (quinpirole)-induced increase in locomotion is blunted in n-3 PUFA deficient animals, consistent with an impairment in β-arrestin2 recruitment and downstream signalling. Yet, however, we find that aripiprazole-induced decrease in motivation – which might result from an antagonistic activity of aripiprazole at the D2R - is also blunted in n-3 PUFA deficient animals. D2R-mediated modulation of motivation has been shown to be independent of β-arrestin2-dependent signalling ^64^, but rather to rely on G_i/o_ protein-mediated transmission at synaptic terminals in the ventral pallidum ^65^. This raises the intriguing hypothesis that membrane PUFAs could differentially modulate D2R signalling activity depending on neuronal subcompartments. Even though further work will be needed to disentangle the precise mechanisms by which PUFAs modulate D2R activity *in vivo*, these latter data are in line with the recent demonstration that motivational deficits in n-3 PUFA deficient animals directly relates to a dysfunction of D2R-expressing neurons ^14^.

Unveiling the precise mechanisms by which brain PUFA biostatus impacts D2R-dependent signaling and downstream-related behaviors will require further work. Nonetheless, taken together, our results show that n-3 PUFAs could act as allosteric modulators of the D2R and potentiate its signalling activity, in particular the β-arrestin2 component. This raises the intriguing hypothesis that n-3 PUFA supplementation could allow potentiating the efficacy of D2R ligands, several of them being antipsychotics. In accordance with this idea, several clinical trials have shown that n-3 PUFA supplementation accelerates treatment response, improves the tolerability of antipsychotics in first-episode psychoses ^66,67^, and reduces prescription rate of antipsychotics ^68^. Our data suggest that direct action of PUFA membrane composition on D2R activity could mediate such alleviating effects.

## Supplementary figures

**Figure S1.**
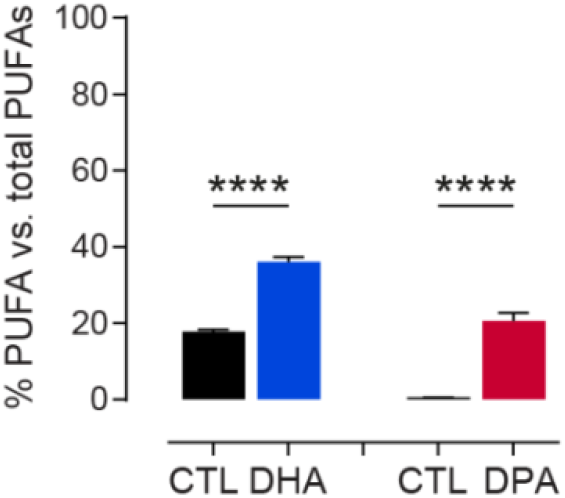
PUFAs enrichment in HEK cells. HEK cells were incubated with DHA or DPA at 10 μM concentration. Each PUFA enrichment is compared to control cells incubated with vehicle only (0.03 % ethanol). Bars represent the mean average of three independent experiments and error bars represent the standard error of the mean (SEM). **** p < 0.0001 by two-tailed unpaired t-test.

**Figure S2.**
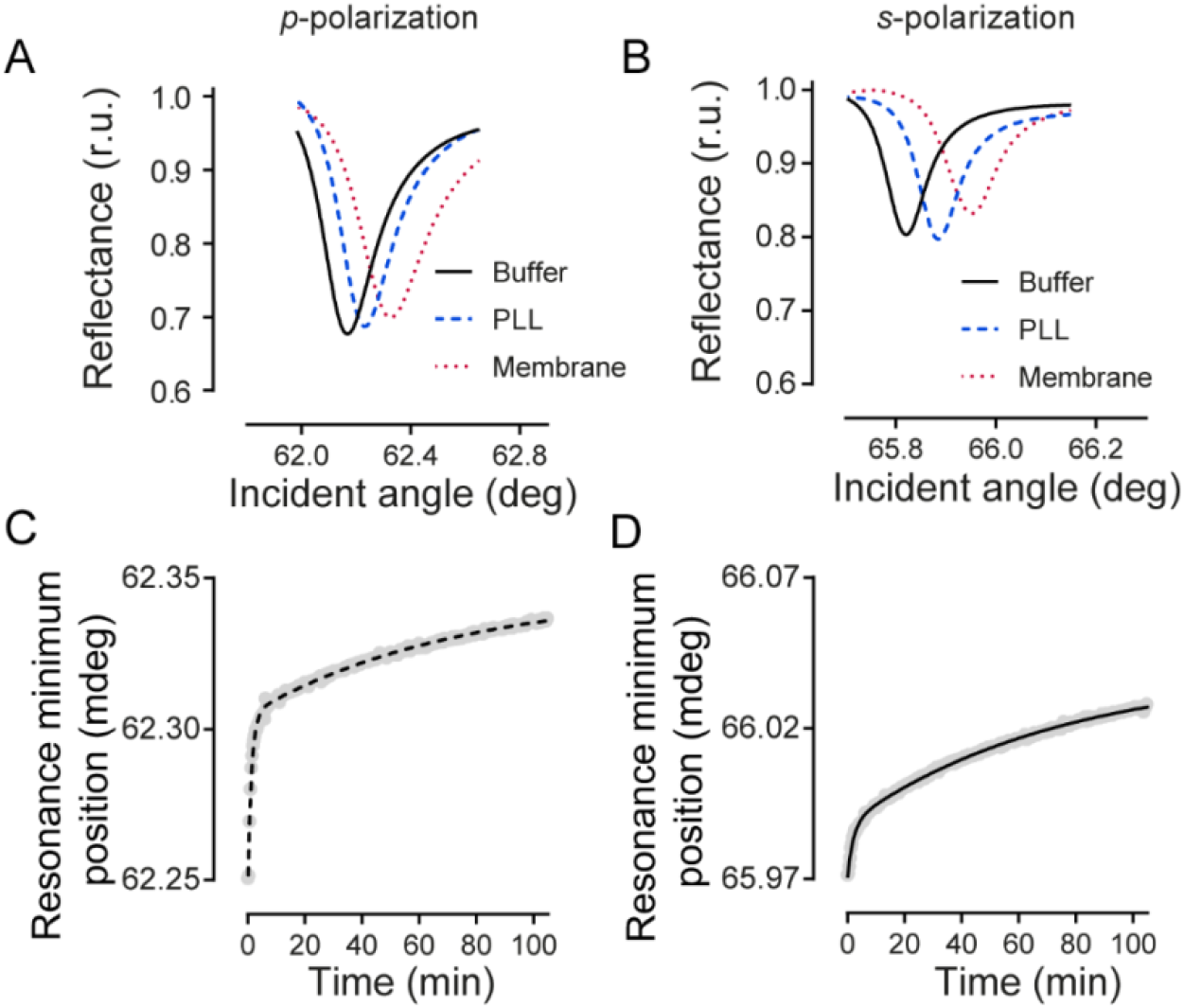
Capture of cell membrane fragments on the PWR sensor and reconstitution of detergent-solubilised D2R in a lipid model membrane. Capture of cell membrane fragments containing the D2R following PLL treatment of the PWR sensor followed by PWR using *p*- and *s*- polarization. (**A and B)** Representative PWR spectra following sensor coating with polylysine (PLL; blue) and cell membrane fragment capture (red) obtained with *p*- (A) and *s*- (B) polarized light, respectively. **(C and D)** Kinetics of D2R reconstitution in a POPC lipid membrane observed for *p*- (C) and *s*-polarizations (D). Kinetics were fitted to a two-phase exponential growth.

**Figure S3.**
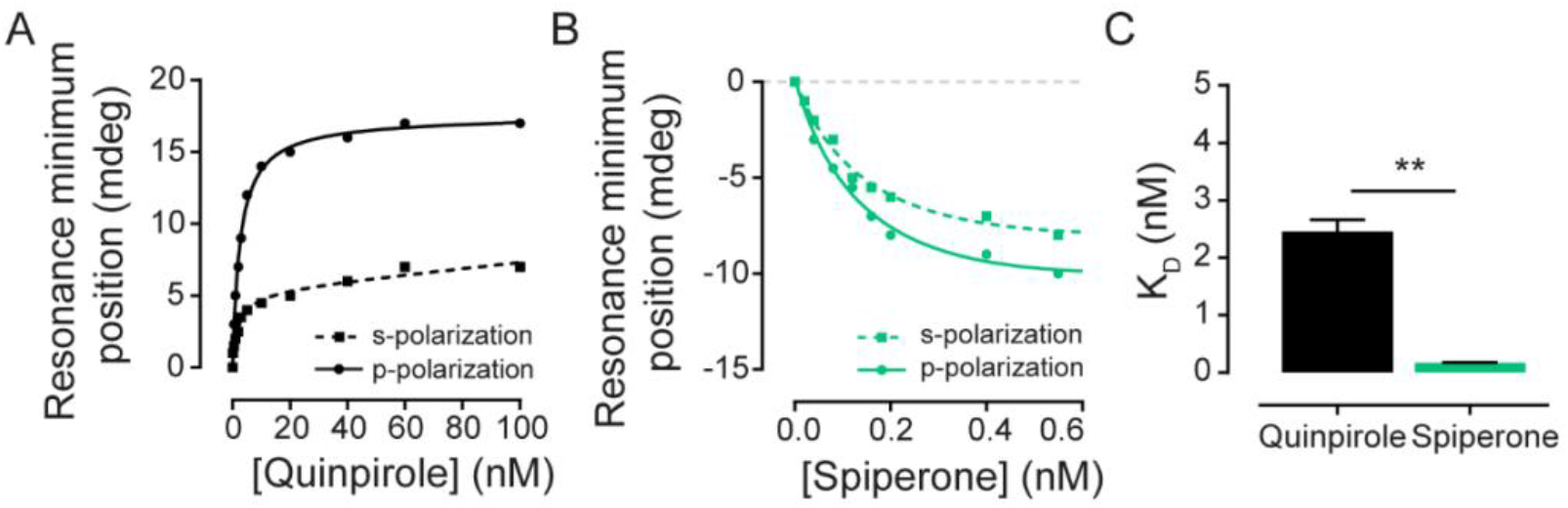
Agonist and antagonist binding with D2R present in cell membranes monitored by PWR. Quinpirole **(A)** and spiperone **(B)** were incrementally added to the proteolipid membrane and the shifts in the resonance minimum position followed. The data were fitted with a hyperbolic binding equation that describes total binding to a single site in the receptor (more details in Materials and Methods). **(C)** Affinity binding of quinpirole and spiperone calculated from **(A)** and **(B)** respectively. ** p < 0.01 by two-tailed unpaired t-test.

**Figure S4.**
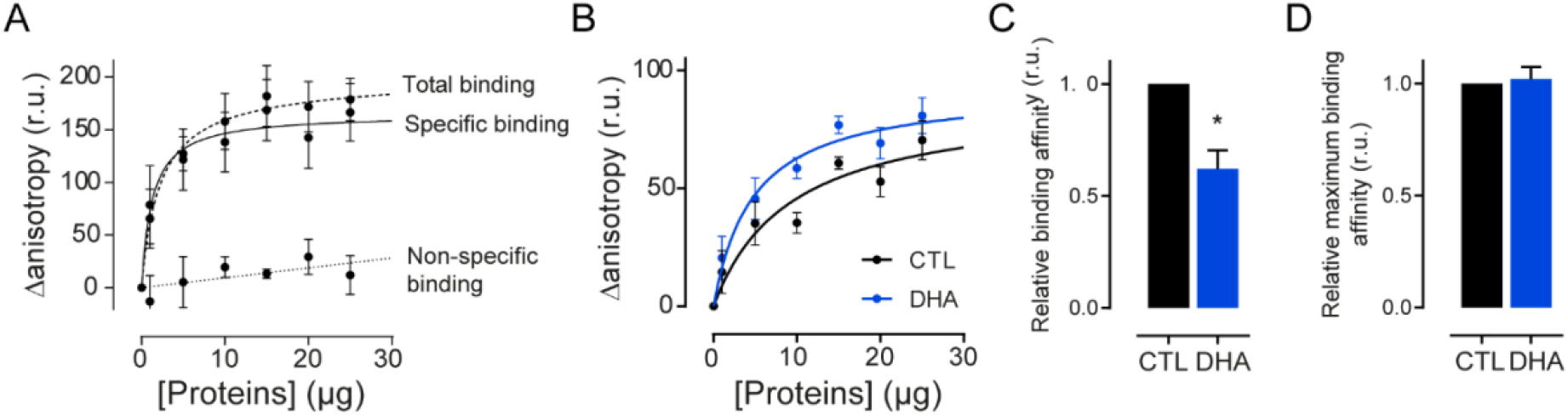
Impact of membrane DHA on D2R ligand binding measured by fluorescence anisotropy. **(A)** Fluorescence anisotropy measurement of NAPS-d2 binding on D2R-expressing membranes (total binding) or non-expressing membranes (non-specific binding). **(B)** Antagonist (NAPS-d2) binding to control and DHA-enriched membranes. **(C)** Fold change of ligand affinity to D2R in DHA-enriched membranes compared to control membranes. **(D)** Maximum relative binding affinity to D2R in DHA-enriched membranes. Data are mean ± SEM from three independent experiments with * p<0.05 by two-tailed unpaired t-test.

**Figure S5.**
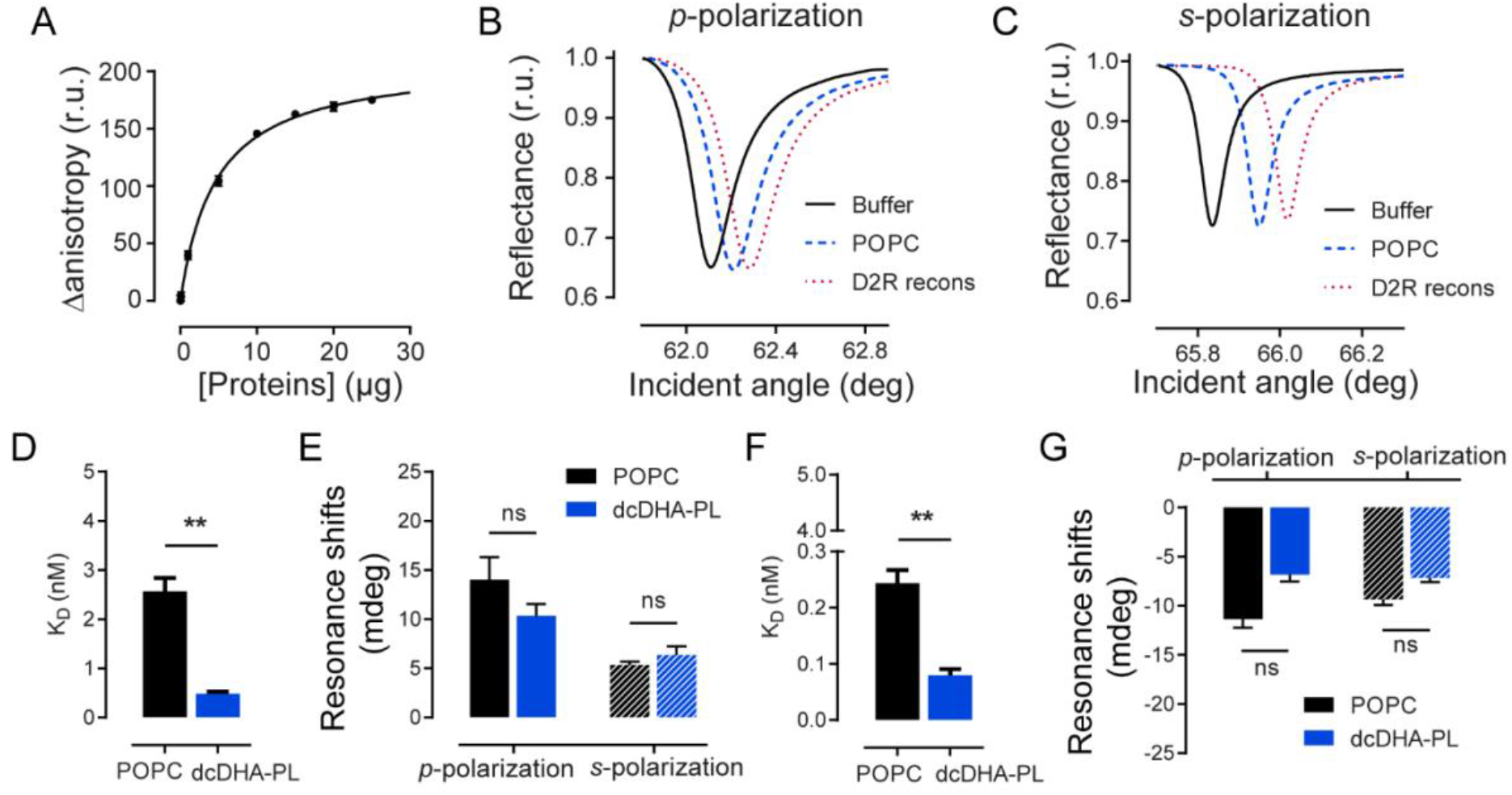
Impact of DHA on ligand binding to D2R partially purified and reconstituted in model membranes shown by PWR. **(A)** Fluorescence anisotropy measurement of NAPS-d2 binding to partially purified D2R (total binding). **(B and C)** Representative PWR spectra following supported POPC lipid membrane formation (blue) and D2R membrane reconstitution (red) for *p*- (B) and *s*-polarized (C) light, respectively. **(D and E)** Effect of DHA on quinpirole affinity to D2R **(D)** and quinpirole-induced receptor conformational changes **(E)** in reconstituted lipid model systems. **(F and G)** Effect of DHA on spiperone affinity to D2R **(F)** and spiperone-induced receptor conformational changes **(G)** in reconstituted lipid model systems. Data are mean ± SEM values from at least three independent experiments with ** p<0.01 by two-tailed unpaired t-test.

**Figure S6.**
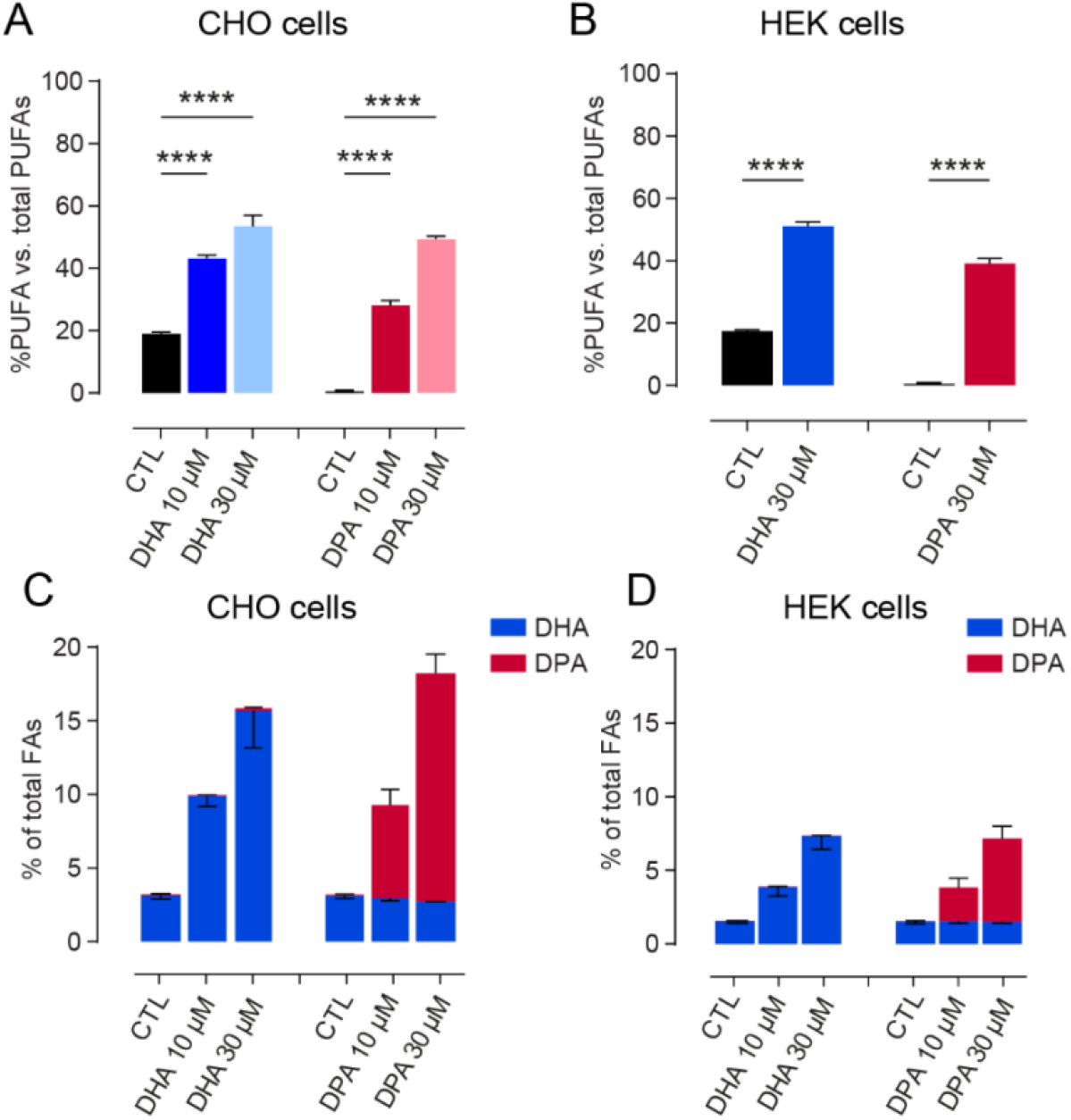
PUFAs enrichment in CHO and HEK cells. **(A)** DHA and DPA incorporation in CHO cells through incubations at 10μM or 30μM. **(B)** DHA incorporation in HEK cells through incubations at 30μM. (C) Cumulative PUFA incorporation in CHO cells through incubations at 10μM or 30μM. (D) Cumulative PUFA incorporation in HEK cells through incubations at 10μM or 30μM. Bars represent the mean average of three independent experiments and error bars represent the standard error of the mean (SEM). ** p < 0.01, *** p < 0.001, **** p < 0.0001 by two-tailed unpaired t-test.

**Figure S7.**
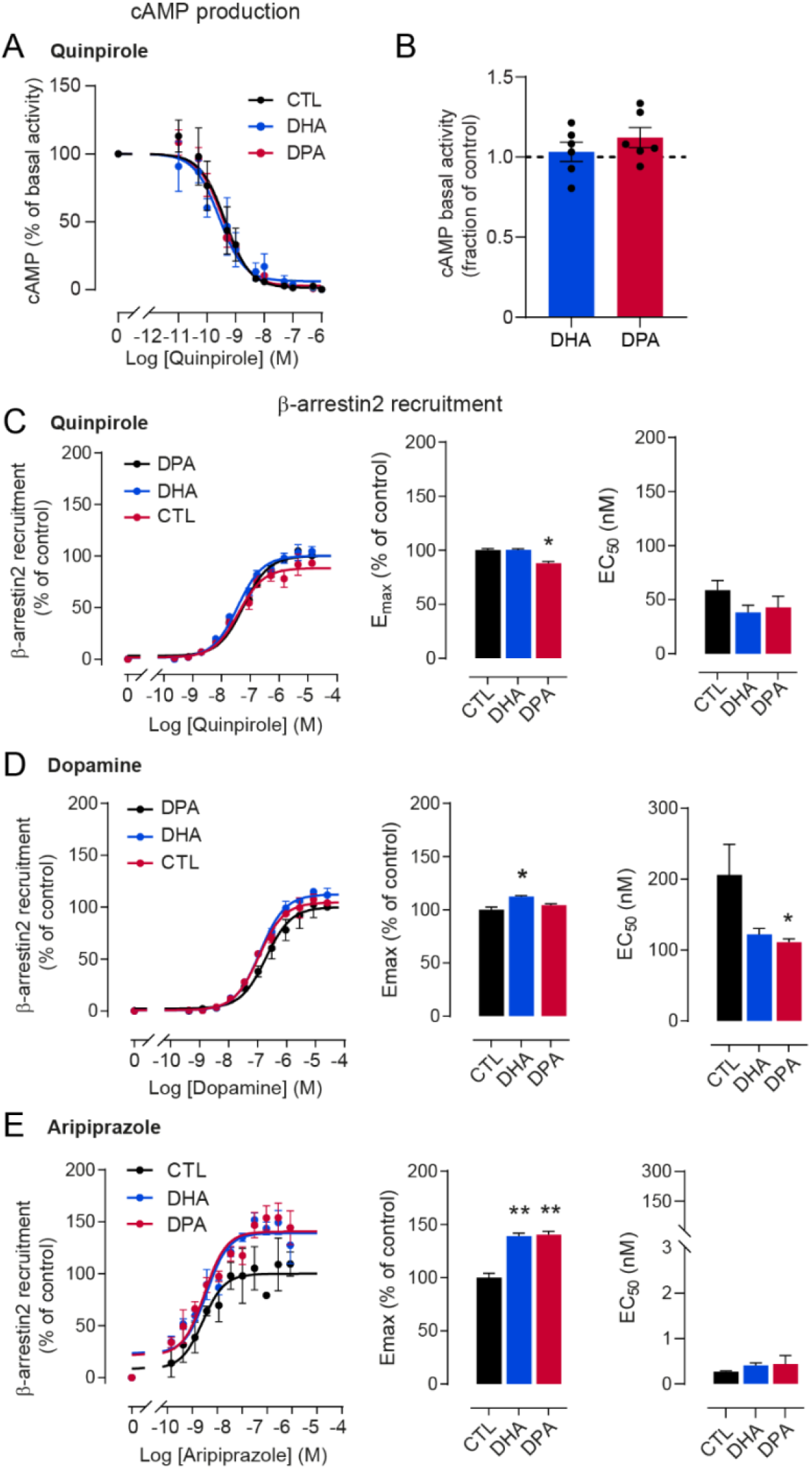
β-arrestin2 recruitment of D2R upon stimulation by different ligands in cells incubated in the presence of vehicle, 10 μM DHA or 10 μM DPA. **(A)** Dose-response experiments of cAMP production with the D2R ligand quinpirole on forskolin-stimulated HEK cells incubated in the presence of 0.03% ethanol as control, 10 μM DHA and 10 μM DPA n-6. **(B)** Effect of 30 μM PUFA enrichment on basal forskolin-induced cAMP production. **(C)** Quinpirole activity on β-arrestin2 recruitment at the D2R in CHO-K1 cells expressing the DRD2L (left), and associated E_max_ (middle) and EC_50_ (right). **(D)** Dopamine activity on β-arrestin2 recruitment at the D2R in CHO-K1 cells expressing the DRD2L (left), and associated E_max_ (middle) and EC_50_ (right). **(E)** Aripiprazole activity on D2R mediated β-arrestin2 assay in CHO-K1 cells expressing the DRD2L (left), and associated E_max_ (middle) and EC_50_ (right). * p < 0.05, ** p < 0.01 by two-tailed unpaired t-test.

**Figure S8.**
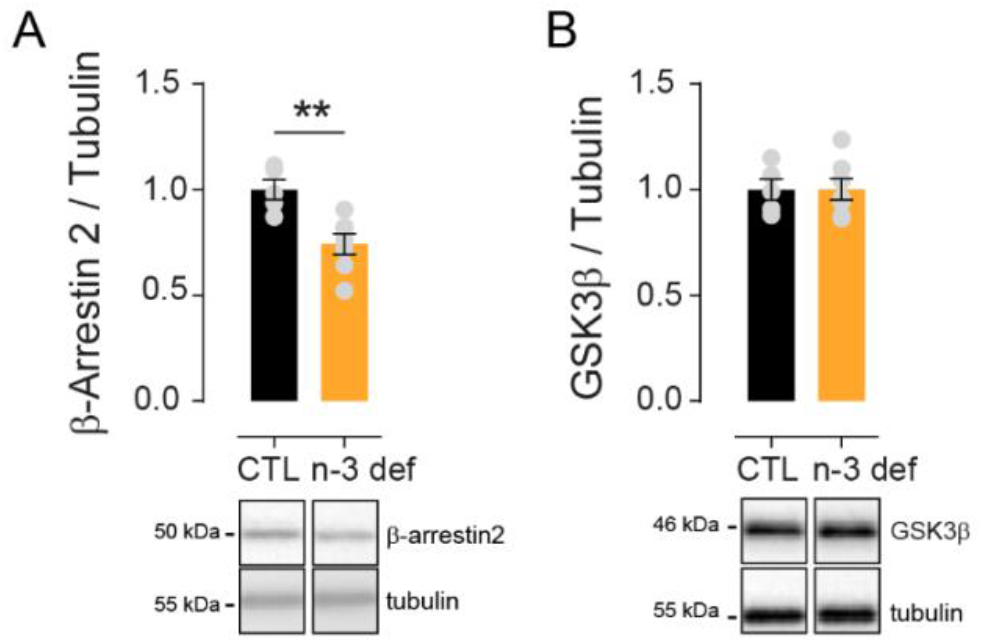
Expression of βarrestin-2 and GSK3β in n-3 deficient mice (n-3 def). **(A) and (B)** Expression of βarrestin2 (A) and GSK3β (B) by western blot normalized to the intensity of tubulin expression. Bars represent the mean of five and seven subjects for CTL and n-3 def respectively and error bars the SEM. **p<0.01 by Mann Whitney test.

**Table S1.** pIC_50_ values (as mean ± SEM) from dose-response experiments of cAMP production on forskolin-stimulated cells incubated in the presence of 0.03% ethanol as control (CTL), 10 μM DHA and 10 μM DPA or 30 μM DHA and 30 μM DPA upon quinpirole, dopamine and aripiprazole stimulation of the D2R.

|  | *CTL* | *DHA 10μM* | *DPA 10μM* | *CTL* | *DHA 30μM* | *DPA 30μM* |
| --- | --- | --- | --- | --- | --- | --- |
| **Quinpirole** | 9.50 ± 0.12 | 9.39 ± 0.25 | 9.48 ± 0.20 | 9.40 ± 0.18 | 9.51 ± 0.08 | 9.62 ± 0.20 |
| **Dopamine** | - | - | - | 8.70 ± 0.12 | 8.54 ± 0.20 | 8.85 ± 0.21 |
| **Aripiprazole** | - | - | - | 8.08 ± 0.25 | 8.36 ± 0.16 | 8.46 ± 0.10 |

**Table S2.**
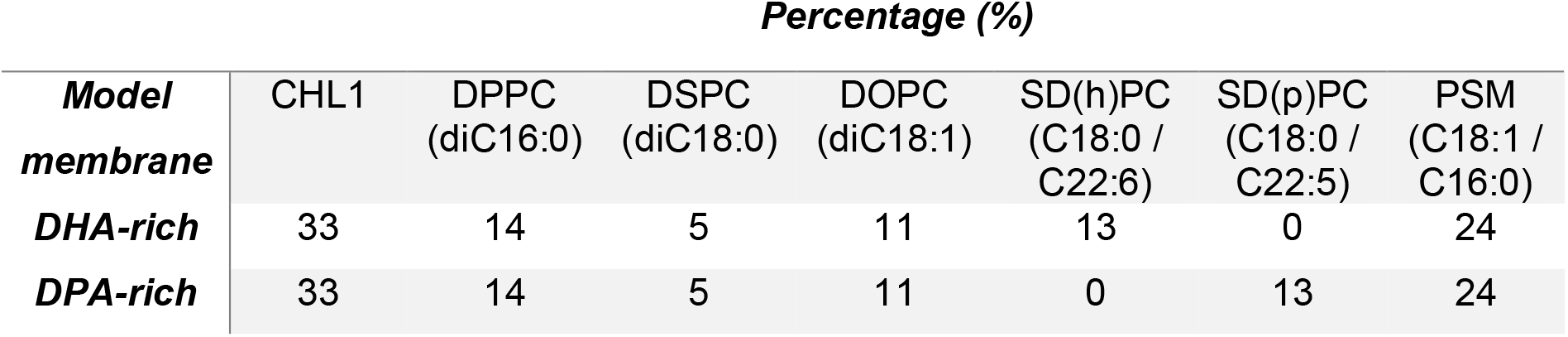
Lipid composition of the model membranes used in MD simulations. CHL1: cholesterol, DPPC: 1,2-dipalmitoyl-sn-glycero-3-phosphocholine, DSPC: 1,2-distearoyl-sn-glycero-3-phosphocholine, DOPC: 1,2-dioleyl-sn-glycero-3-phosphocholine, SD(h)PC: 1-stearoyl-2-docosahexaenoyl-sn-glycero-3-phosphocholine, SD(p)PC: 1-stearoyl-2-docosa(p)entaenoyl -sn-glycero-3-phosphocholine, PSM: sphingomyelin.

## Material and Methods

### Chemicals and antibodies

Reagents, unless otherwise specified, were from Sigma-Aldrich (St. Louis, MO, USA) PUFAs (Cis-4,7,10,13,16,19-DHA (DocosaHexaenoic Acid, ref D2534), DPA n-6 (DocosaPentanoic Acid, ref 18566) stock solutions (30mM) were prepared in absolute ethanol under N2 (De Smedt-Peyrusse et al., 2008). For anisotropy assays, LigandTag Lite D2 (L0002RED) receptor red antagonist was purchased from Cisbio Bioassays. Quinpirole (ref 1061) and Forskolin (ref 1099) was from Tocris.

### Cell culture and treatment

HEK 293 stably expressing the human D2R (Flp-in T-rex 293 SF-D2sWT/FRTTO 293) were used. Cells were maintained at 37°C under 5% CO2 in the following culture medium: DMEM (Dulbecco’s modified Eagle’s medium) Glutamax (InVitroGen) containing 10% of heat inactivated fetal bovine serum, 100UI/ml penicillin and 0.1mg/ml streptomycin and supplemented with 100μg/ml Hygromycin (InVitrogen), 15μg/ml Blasticidin (Cayla Invivogen) and 1μg/ml Tetracycline (Sigma Aldrich). Upon reaching 80-90% confluence, in the same culture medium, cells were treated with 10μM and 30 μM DHA or DPA or 0.03% and 0.01% ethanol (vehicle) respectively as control for 24h. After treatment, cells were dissociated by using an Enzyme-Free Cell Dissociation Solution (S-014-B, Merck Millipore) and washed twice in wash buffer (50mM Tris-HCl, 50mM NaCl, 2mM EDTA, pH 7.5 at 20°C) by centrifugation at 500 x g for 8 min at 20°C. Cell pellets were used either for Western blot analyses or for membrane preparation.

### Membrane preparation

The cell pellets of HEK cells were resuspended in 10 ml of lysis buffer (10 mM Tris-HCl, 1 mM EDTA, pH 7.5 at 4°C) supplemented with anti-protease inhibitor cocktail (PIC) (P8340, Sigma Aldrich) and incubated for 30 min in ice. The cell lysate was homogenized using a 10 ml glass-Teflon homogenizer with 30 gentle strokes. The resulting suspension was centrifuged at 183 x g for 10 min at 4°C. The supernatant was collected and the residual pellet was washed twice by centrifugation in successively in 5 and 2.5ml of lysis buffer. The resulting supernatants were pooled and centrifuged at 48000 x g for 30 min at 4°C. The pellet containing the membrane proteins was resuspended in binding buffer (50 mM Tris-HCl, 50 mM NaCl, 2 mM EDTA, 2 mM CaCl_2_, pH 7.5 at 4°C) supplemented with PIC. Protein concentration was determined by using Bicinchoninic Acid (BCA) protein assay (Uptima, Montluçon, France) and membranes were diluted to 1μg/μl in the binding buffer.

### Plasmon waveguide resonance (PWR)

PWR experiments were performed with a homemade instrument equipped with a He-Ne laser at 632 nm whose light is linearly polarized at 45°, allowing acquisition of both *p*- (light that is parallel to the incident light) and *s*-polarization (light that is perpendicular to the incident light) data within a single angular scan. The technique has been described previously ^30,69^. Experiments were carried out at controlled room temperature, e.g. 23°C. The sensor consists in a 90° angle prism whose hypotenuse is coated with a silver layer (50 nm) and overcoated with silica (460 nm) and is in contact with the cell sample Teflon block, with an aperture of approximately 3 mm diameter through which the lipid bilayer is formed. This is placed on a rotating table mounted on a corresponding motion controller (Newport, Motion controller XPS; ≤ 1 mdeg resolution).

### Formation of a planar lipid bilayer on the PWR sensor

The method used to prepare the lipid bilayer is based on the procedure by Mueller and Rudin ^70^ to make black lipid membranes. The planar lipid bilayer is formed across the small aperture (~ 3 mm) in the Teflon PWR cell by fusing freshly sonicated small unilamellar vesicles (SUV, 3mg/ml) with the silica surface. SUVs were prepared by initially dissolving the appropriate amount of phospholipids in chloroform, to obtain the desired final concentration. A lipid film was then formed by removing the solvent using a stream of N2 (g) followed by 3h vacuum. The lipid film was dispersed in PBS and thoroughly vortexed to form multi-lamellar vesicles (MLVs). To form SUVs, the MLVs dispersion was sonicated 5 times during 10 minutes at 40Hz frequency just before use. To ensure that a proper solid-supported lipid bilayer is formed, the changes in the resonance minimum position (resulting from changes in mass density, anisotropy and thickness following film deposition) for both polarizations are measured and compared to values previously established to correspond to a lipid bilayer ^25^.

### Immobilisation of cell fragments on the PWR sensor

The protocol for adhesion of cell fragments on silica (glass slides or PWR sensor) was adapted from reported work from Perez and collaborators ^71^. Briefly, the silica surface was washed with ethanol, cleaned and activated by Plasma cleaner for 2 min (Diener, Bielefeld, Germany). The silica surfaces were then incubated with a solution of poly-L-lysine (PLL, 0.1 mg/mL) for 40 minutes following wash with PBS buffer. Cells grown to less than 50% confluence were washed with PBS and covered with water to induce osmotic swelling of the cells. Immediately, the glass coverslip of the sensor was placed directly on top of cells. Pressure was applied for about 2 min on the glass slide or prism to induce cell rupture and caption of cell fragments. Then they were removed ripping off cell fragments containing specially the upper membrane. The glass slide or sensor was washed with PBS to remove cell debris and kept with buffer to prevent drying and loss of membrane protein activity. PWR measurements were performed right away. The PWR cell sample (volume capacity of 250 μL) was placed in contact with the prism and filled with PBS.

### Reconstitution of the dopamine D2 receptor in the lipid bilayer

After lipid bilayer formation, detergent-solubilized D2 receptor was reconstituted in the lipid bilayer by the detergent-dilution method. Briefly, the receptor was purified in a mixture of dodecylmatoside and cholesteryl hemisuccinate (DDM/CHS) at concentrations that are about 10 fold over the critical micelle concentration (cmc) of DDM. Part of the detergent was then removed from the sample by use of centricons (Merck Millipore) with a cutoff of 50 KDa. This consisted in concentrating and reducing the initial volume of the solubilized protein by a factor of 5. The insertion of a small volume (about 20 μL) of DDM/CHS/OG solubilized protein into the PWR chamber, leads to drastic and quick drop in the detergent concentration. If the detergent concentration drops below the cmc during this step, this results in immediate positive PWR shifts for both *p*- and *s*-polarisations with small changes in the TIR angle. In order to compare data among different experiments, data were normalized relative to the amount of reconstituted protein in the membrane.

### Ligand-induced receptor response

Both systems (cell fragments and reconstituted protein in lipid model systems) were tested for their capacity to respond to ligand to check if the dopamine D2 receptor is active after cell fragment immobilisation or reconstitution process and if it was properly reconstituted in the lipid membrane. Briefly, the method consisted in incrementally adding a specific dopamine D2 ligand to the cell and monitor the PWR spectral changes. The first concentration point was chosen to be approximately 1 order of magnitude lower than the published dissociation constant (K_D_) value for that ligand. Before each incremental concentration of ligand added, the system was left to equilibrate. K_D_ values were obtained from plotting the resonance minimum position of the PWR spectra (this reflects the receptor-ligand complex) as a function of total ligand concentration and fitting to the hyperbolic function that describes the 1:1 binding of a ligand to a receptor using GraphPad Prism (GraphPad Software).

A control experiment was performed that consisted in adding the same ligand concentrations to a lipid bilayer with no receptor reconstituted. This measured non-specific binding of ligand to lipids alone (data not shown).

### Fluorescence anisotropy assay

Fluorescence anisotropy (FA) was used to measure receptor-ligand binding reaction into individual wells of black, 96-well Greiner Bio-One microplates in a final volume of 100μl. To establish a saturation binding curve, the range of protein membrane concentration used was from 1 to 25μg in the presence of 10nM of the antagonist NAPS-d2 ligand (L0002 red, Cisbio Bioassays). The FA was measured on a Tecan Infinite M1000 Pro microplate reader (Männedorf, Switzerland). Excitation was set at 590 ± 5 nm and emission was collected at 665 ± 5 nm bandpass filters for the Texas Red.

The non-specific binding was obtained from membranes that did not express the receptor and the total binding was measured with D2R-expressing membranes. The non-specific binding curve was fitted with a Simple-linear regression in GraphPad Prism (GraphPad Software, San Diego, CA), the total binding curve was fitted with a One site – total binding curve in GraphPad Prism. The specific binding was determined by subtracting the non-specific curve to the total binding curve and the final specific binding curve was fitted with a One site – specific binding model in GraphPad Prism. The latter was used to determine relative K_D_ values via separated saturation binding curves in at least three independent experiments and are reported as mean ± SEM.

### cAMP accumulation assays

D2R stably expressing HEK 293 cells were grown on PLL treated 12-well plates as described above. Upon reaching 80-90% confluence, cells were treated with 10 μM or 30 μM DHA or DPA n-6 or 0.03 % to 0.1 % ethanol (vehicle) as control for 24 h in the culture medium. The medium was removed and cells were rinsed in DMEM before pretreatment with 1mM IsoButylMethylXanthine (IBMX, Sigma I5879) for 15min. Cells were then stimulated for 30min with the indicated concentrations of agonists Quinpirole (Tocris 1061), Dopamine (Sigma H8502) and Aripiprazole (Sigma SML0935) in the presence of 1mM IBMX and 10μM Forskolin (Tocris 1099). Endogenous phosphodiesterase activity was stopped by removing the medium and the addition of 0.1M HCL (300μl/well). After centrifugation at 600g during 10min, protein concentration of supernatants was quantified by BCA. Cyclic AMP levels were determined in samples containing 10μg of protein. The production of cAMP was measured by using a cAMP Enzyme Immunoassay kit (Sigma, CA200) as described by the manufacturer using Victor3 (Perkin Elmer) plate reader. The curve fit was obtained by GraphPad Prism 5 (GraphPad Software, Inc.).

### β-Arrestin2 recruitment assay

β-arrestin2 recruitment was assessed using the PathHunter^©^ express DRD2L CHO-K1 Beta arrestin GCPR Assay (DiscoverX, Fremont, CA). In brief, cells were plated into 96-well white-walled assay plates in a volume of 90μl of Cell Plating Reagent (DiscoverX). They were incubated 24 h at 37 °C, 5 % CO_2_.The next day, PUFAs (Ethanol 0.01% as control vehicle, DHA, and DPA n-6) were prepared at 10X concentration (300μM) in Cell Plating Reagent and 10 μL was added to the cells for 24h at 37 °C, 5 % CO_2_. Serial dilutions (11x) ranging from 154 to 0.0026 μM, 281 to 0.0046 μM and 9.35 to 0.000157 μM, of Quinpirole, Dopamine and Aripiprazole respectively, were prepared and 10 μl of each concentration was added for 90 minutes. Luminescence was measured at 1 h post PathHunter^©^ detection reagent addition using Victor3 plate reader (Perkin Elmer 0.5-s/well integration time). Data were normalized to control treated with the highest concentration of ligand (100%). This control value corresponds to Top best fit value determined by non-linear fit of standard slope. Data were fitted to a three-parameter logistic curve to generate EC_50_ and E_*max*_ values (Prism, version 5.0, GraphPad Software, Inc., San Diego, CA). EC_50_ and % E_max_ values are the result of three independent experiments performed in duplicate.

### Animals and experimental model

Mice were housed in groups of 5-10 animals in standard polypropylene cages and maintained in a temperature and humidity-controlled facility under a 12:12 light-dark cycle (8:00 on) with ad libitum access to water and food. All animal care and experimental procedures were in accordance with the INRA Quality Reference System and to french legislations (Directive 87/148, Ministère de l’Agriculture et de la Pêche) and European (Directive 86/609/EEC). They followed ethical protocols approved by the Region Aquitaine Veterinary Services (Direction Départementale de la Protection des Animaux, approval ID: B33-063-920) and by the animal ethic committee of Bordeaux CEEA50. Every effort was made to minimize suffering and reduce the number of animals used. C57BL/6J mouse lines from Janvier Laboratories (Robert Janvier, Le Genest St-Isle France) were used in this study. Female C57BL6/J mice were fed with isocaloric diets containing 5% fat with a high (n-3 def diet) or low LA/ALA ratio (Ctrl diet) across gestation and lactation and offspring were maintained under the same diet after weaning as previously done (see details in ^14^). All experiments were performed at adulthood.

### Operant conditioning

The apparatus used for these experiments was previously described ^14^. Animals were food-restricted in order to maintain them at 85-90% of their ad libitum weight and exposed to one session (1 hour) each day, 5-7 days per week. A pavlovian training followed by fixed and random ratio training were performed in order to make the animals reach an acquisition criterion (i.e. the number of lever presses and rewards earned) before performing the motivational task per se as previously described ^14^. In the motivational task, namely the progressive ratio times 2 (PRx2) schedule, the number of lever presses required to earn a reward was doubled respective to the previous one obtained. Mice were tested multiple times in PRx2 with RR20 sessions intercalated between each PR tasks. Aripiprazole (Arip; 7-[4-[4-(2,3-dichlorophenyl)piperazin-1-yl]butoxy]-3,4-dihydro-1H-quinolin-2-one; Merck®, Darmstadt, Germany) was dissolved in a mixture of saline (NaCl 0.9%) and cremiphore (2% of cremiphore in saline) at the doses of 0.1 and 0.5 mg/kg and administered intraperitoneally 10 min before the beginning of the PRx2 test sessions as previously done (Ducrocq et al., in prep.). The ratio of lever presses under drugs over lever presses after vehicle injection were measured for each animal and averaged in order to evaluate the effect of the drugs on operant responding.

### Spontaneous locomotion

Animals were transferred individually to small Plexiglas cages (10 cm wide, 20 cm deep, 12 cm tall) equipped with a video tracking system (Smart, Panlab, Barcelona, Spain) allowing the recording of total distance travelled (cm) as a measure of spontaneous locomotor activity in basal condition for one hour. Quinpirole-induced locomotor response was evaluated as described previously (Akhisaroglu et al., 2005). Briefly, animals received 7 intraperitoneal injection of 1mg/kg quinpirole (Quin; Tocris) dissolved in 0.9% saline every 72 hours. Locomotion was immediately measured for 3 hours after the 8th injection of 0.5mg/kg Quin.

### Western blot

Mice were dislocated and the brains were quickly removed, snap-frozen on dry ice and stored. Nucleus accumbens samples were punched (No.18035-01, Fine Science Tools) from 200 μm frozen slices in a cryostat. Samples were homogenized and denatured and western blot were performed as previously described ^14^. Briefly, equal quantities of proteins (10μg/well) were separated by electrophoresis. Membranes were saturated and blots were probed with the following primary antibodies: 1:700 rabbit anti-βArr2 (Cell signaling Cat. No. 3857); 1:1000 rabbit anti-GSK3β (Cell signaling Cat. No. 12456); 1:1000 rabbit anti-P-GSK3β (Cell signaling Cat. No. 5558); 1:10000 mouse anti-tubulin (Merck Cat. No. T5168). Then, membranes were incubated with the secondary antibody coupled to Horse Radish Peroxidase (HRP, 1/5000, Jackson ImmunoResearch). After revelation, optical density capture of the signal was performed with the ChemidocMP imaging system (BioRad) and intensity of the signal was quantified using the Image Lab software (BioRad).

### Cell Membrane Microarray and [^35^S]GTPγS autoradiography

Microarrays were composed of a collection of membrane homogenates isolated from the NAc of adult mice exposed to Ctrl diet (n=17) or n-3 def diet (n=17) and from rat cerebral cortex as positive control. Briefly, tissue samples were homogenized using a Teflon-glass grinder (Heidolph RZR 2020) and a disperser (Ultra-Turrax® T10 basic, IKA) in 20 volumes of homogenized buffer (1 mM EGTA, 3 mM MgCl2, and 50 mM Tris-HCl, pH 7.4) supplemented with 250 mM sucrose. The crude homogenate was subjected to a 3,000 rpm centrifugation (AllegraTM X 22R centrifuge, Beckman Coulter) for 5 min at 4°C, and the resultant supernatant was centrifuged at 14,000 rpm (Microfuge® 22R centrifuge, Beckman Coulter) for 15 min (4 °C). The pellet was washed in 20 volumes of homogenized buffer and re-centrifuged under the same conditions. The homogenate aliquots were stored at −80 °C until they were used. Protein concentration was measured by the Bradford method and adjusted to the required concentrations. Microarrays were fabricated by a non-contact microarrayer (Nano_plotter NP2.1) placing the cell membrane homogenates (20 drops/spot) into microscope glass slides treated using a proprietary technology, which enables the immobilization of cell membranes to supports preserving the structure and functionality of their proteins ^72^.

[^35^S]GTPγS binding studies were carried out using Cell Membrane Microarrays according to the following protocol. Briefly, Cell Membrane Microarrays were dried 20 min at room temperature (r.t.), then they were incubated in assay buffer (50 mM Tris-Cl; 1 mM EGTA; 3 mM MgCl_2_; 100 mM NaCl; 0,5% BSA; pH 7,4) in the presence or absence of 50 μM GDP and/or 100 μM quinpirole for 15 min at r.t.. Microarrays were transferred into assay buffer containing 50 μM GDP and 0.1 nM [^35^S]GTPγS, with and without the dopamine D2 agonist, quinpirole, at 100 μM and incubated at 30°C for 30 min. Non-specific binding was determined with GTPγS (10 μM). Finally, microarrays, together with [^35^S]-standards, were exposed to films, developed, scanned and quantified using the Mapix software.

### Lipid analyses

Cells submitted to different treatments were randomly analyzed according to Joffre et al. ^73^. Total lipids were extracted according to the method developed by Folch et al. ^74^ and were submitted to fatty acid methylation using 7% boron trifluoride in methanol following Morrison and Smith protocol ^75^. Gas chromatography on a HewlettPackard Model 5890 gas chromatograph (Palo Alto, CA, USA) was employed to analyze fatty acid methylesters (FAMEs) using aCPSIL-88 column (100 m×0.25 mm internal diameter;film thickness,0.20 μm; Varian, Les Ulis, France). Hydrogen was used as a carrier gas (inlet pressure, 210 kPa). The oven temperature was maintained at 60 °C for 5 min, then increased to 165 °C at 15 °C/min and held for 1 min, and then to 225 °C at 2 °C/min and finally held at 225 °C for 17 min. The injector and the detector were maintained at 250 °C and 280 °C, respectively. FAMEs were identified by comparison with commercial and synthetic standards and the data were computed using the Galaxie software (Varian). The proportion of each fatty acid was expressed as a percentage of total fatty acids to allow the comparison of lipid composition in different cell culture conditions.

### MD simulations

VMD1.9.4 ^76^ was used to preprocess both inactive crystal (PDB ID: 6CM4) and active cryoEM structures (PDB ID: 6VMS) of the dopamine D2 to set up the simulations of the apo and holo states, respectively. Any co-crystallization atoms different than water molecules closer than 5 Å to the protein were removed. MODELLER ^77^ was used to: (a) mutate back to the native sequence any mutation present in the structure, namely A122I, A375L, and A379L in PDB ID: 6CM4, and I205T, L222R, L374M, Y378V, L381V, and I421V in PDB ID: 6VMS, and (b) model residue CYS443, which is missing in the 6CM4 structure, to have the D2R palmitoylation site available. The HomolWat server ^78^ was used to place internal water molecules not present in the initial structure, and the sodium ion in the case of apo structures. An inactive apo structure was then generated by simply removing the risperidone ligand bound to PDB ID: 6CM4. Likewise, an active apo structure was generated by removing the ligand bromoergocryptine from PDB ID: 6VMS. This second apo structure was used to dock one molecule of either dopamine or aripiprazole into the orthosteric binding pocket of the receptor using AutoDock Vina ^79^.

The CHARMM-GUI builder ^80,81^ was used to embed each refined structure were into a 90 × 90 Å^2^ multicomponent membrane rich in either DHA or DPA, using a specific and realistic lipid composition ^20^, which is detailed in Table S2.

Each protein-membrane system was placed into a water box made of explicit water molecules, their charge was neutralized, and the ionic strength of the system adjusted, throughout CHARMM-GUI builder’s pipeline. All titratable residues of the protein were left in their dominant protonation state at pH 7.0, except for Asp80. Disulfide bridges were inserted between Cys107-Cys182, and Cys399-Cys401, and a palmitoyl moiety was covalently linked to Cys443. Systems were first energy minimized and then equilibrated for 50 ns at constant pressure (NPT ensemble). The harmonic positional restraints initially applied to all Cα atoms of the protein were gradually released throughout the equilibration phase. Production simulations for each replica were run for 1 μs each, at constant volume (NVT ensemble), 1,013 bar and 310 K. The production simulations of this study yielded an aggregated time of 16 μs (4 systems x 4 replicas x 1 μs). All simulations were run using ACEMD ^82^ in combination with the CHARMM36m force field ^83^. Dopamine and aripiprazole ligand charges were optimized in the ANI-2x mode using the parameterize module of HTMD ^84^. Figures from simulations were rendered using the Tachyon renderer ^85^ and the R ggplot2 library ^86^.

Lipid-protein contact ratios between lipid species were calculated by dividing the number of contacts per atom of one lipid group over the other. Atoms closer than 4.2 Å to the centre of mass of the protein were considered in contact. Only lipid tail atoms were used in these calculations. Each contact value was previously normalized by the total number of atoms in that particular selection. For example, sn-1 SDP(d)C / SAT ratios were calculated as the number of atoms of DHA < 4.2 Å of protein’s centre of mass divided by the total number of SDP(d)C chain atoms in the system. Likewise, SAT is calculated in the former ratio as the number of atoms of any DPPC, DSPC, or PSM chain closer than 4.2 Å of protein’s centre of mass divided by the total number atoms of these tails.

The average lipid occupancy across the simulations was computed using the Python package PyLipID ^87^.

### Statistical analyses

Data are reported as mean ± SEM. Statistical analyses were conducted using Prism 6 software (GraphPad Software, La Jolla, CA, USA). Two-tailed unpaired Student’s t-test or Mann Whitney test were used to assess differences between two groups. Two-way RM ANOVA followed by post-hoc Bonferroni tests were used for repeated measures. Differences were considered significant for p < 0.05.

## Acknowledgements

We thank the biochemistry facility of the Bordeaux Neurocampus for the access to the blot imaging system, JP. Toulmé for use of the fluorescence spectrometer (TECAN), JL Banere for providing D2R expressing *Pichia pastoris* and the Bordeaux Metabolome Facility-MetaboHUB (ANR-11-INBS-0010).

This study was supported by INRAE, CNRS and the University of Bordeaux, Idex Bordeaux “chaire d’installation” (ANR-10-IDEX-03-02) (P.T.), NARSAD Young Investigator Grants from the Brain and Behavior Foundation (P.T.), ANR “SynLip” (ANR-16-CE16-0022) (P.T.), ANR “FrontoFat” (ANR-20-CE14-0020) (P.T.), Region Nouvelle Aquitaine 2014-1R30301-00003023 (P.T.), NIH grant MH54137 (J.A.J), the Instituto de Salud Carlos III FEDER (PI18/00094) and the ERA-NET NEURON & Ministry of Economy, Industry and Competitiveness (AC18/00030) (J.S.), the European Research Network on Signal Transduction (https://ernest-gpcr.eu) (COST Action CA18133) (B.M., R.G-G., J.S.), the Swiss National Science Foundation, grant no. 192780 (R.G-G.).

## Author contributions

M-L.J, V.D.P, R.G-G, I.D.A. and P.T. conceived and supervised the study. J.S., G.B-G., E.M., T.D. and J.A.J. provided expertise, reagents and supervised specific experiments. M-L.J., V.D.P., F.D., A.O., R.B., M.H.P., B.M-L., J.S., S.M., T.T-C., S.G. and R.G-G performed experiments and analyzed the data. M-L.J., V.D.P., R.G.-G., I.D.A. and P.T. wrote the original version of the manuscript. All authors discussed the results and reviewed the manuscript.

